# An Interpretable Multi-instance Learner Decodes Cellular Recruitment from Spatially Resolved Transcriptomics

**DOI:** 10.1101/2025.11.20.689581

**Authors:** Jia Yao, Ji Seon Shim, Kelly Wong, James Zhu, Yunhe Liu, Sung Wook Kang, Kelli Bryan M. Burt, Jong Min Choi, Claire Lee, Idania Carolina Lubo Julio, Luisa Maren Solis Soto, Sharia Hernandez, Karen Colbert, Wei Lu, Khaja Khan, Paul F Mansfield, Hee-Jin Jang, Tao Yue, Changwei Li, Hao Zhu, Linghua Wang, Hyun-Sung Lee, Tao Wang

**Author notes:** Co-first authors (Jia Yao, Ji Seon Shim).

## Abstract

The recruitment of the various types of cells into the tissue microenvironment and how these cells engage with other cells in the tissue sites play critical biological roles. However, it is difficult to study these processes on a genome-wide scale using traditional low-throughput experiments. To address this gap, we developed a fully interpretable multi-instance deep learner, coined “spacer”. Spacer digests the transcriptomics and spatial modalities of spatially resolved transcriptomics (SRT) data in alignment with the biological principles of the tissue spatial architecture. We deployed spacer to a panel of 17 high definition and 20 low definition SRT datasets, for studying how stromal and immune cells were recruited into tumors and heart during myocarditis. Spacer discovered novel biological insights not affordable by prior spatial data analysis tools, which were validated by orthogonal immuno-peptidomics, spatial T cell receptor (TCR) sequencing, and single cell sequencing experiments. We discovered genes that encode more immunogenic peptides and that are involved in developmental pathways are more potent in recruiting T cells to local tumor sites, while the recruitment of B cells and macrophages is stipulated by a different molecular program. On the other hand, expression of mucins in the tumor cells was found to repel T cell localization. For the engaging cell side, we also uncovered T cell-intrinsic features that determine their localization, validated by spatial-TCR-seq data. Spacer also unexpectedly revealed that CD4^+^ T cells, though fewer in numbers, are more responsive than CD8^+^ T cells in the heart during myocarditis. Collectively, this study establishes a new spatially resolved paradigm for studying cellular localization mechanisms *in situ* and shifts the paradigm from descriptive mapping to mechanistic discovery.

## Main

Various cell types are recruited into tissue microenvironments based on intrinsic and extrinsic signals, which enable these cells to execute key biological functions. Extensive research has been performed to investigate these processes, but most biological experiments examine one cell type or pathway at a time, and usually involve *in vitro* artifacts that could result in physiologically irrelevant conclusions. A global unbiased approach is lacking to generate hypotheses regarding cellular recruitment and engagement in unperturbed *in situ* conditions. The recent rise of high definition whole-transcriptome (WTX) spatially resolved transcriptomics (SRT), such as VisiumHD, Slide-Tag ^1^, CosMx WTX, and Stereo-Seq ^2^, pushes the boundary of achieving (near) single cell resolution transcriptomes of more cells. However, SRT data analyses are challenging due to the sheer amount of data and the complicated data structures involving the cells. They have thus remained largely descriptive, namely cataloging what is where, without revealing why cells localize as they do.

To fill this critical void, we introduce an algorithm called Spatial Analysis of Cellular Engagement and Recruitment (spacer), a fully interpretable multi-instance deep learner designed to decrypt the “spatial language” of tissues. Representing a departure from conventional black-box model, spacer pioneers an interpretable multi-instance architecture that digests both the transcriptomics and spatial modalities of SRT data through an attention mechanism. Our central hypothesis is that the rules of cellular recruitment and engagement can be ascertained from spatial patterns. By rendering the decision-making process transparent, spacer advances beyond mere description to revealing human-interpretable biological rules, enabling the discovery of translationally applicable targets directly from real-word clinical biospecimens. To achieve this, we reason that AI models must possess a structure that is isomorphic to the biological tissues. Namely, the structure of the deep learning model should be reflective of the actual biological system, which is the spatial architecture of the tissue in question here. Therefore, spacer embeds the biological principles of cellular neighborhoods directly into its architecture, establishing a new paradigm for mechanistic discovery.

We used spacer to comprehensively characterize the biology of stromal and immune cell recruitment and engagement in human tumors in a pan-cancer manner. We deployed spacer to analyze high definition WTX SRT datasets generated from the VisiumHD, Slide-tag, and CosMx WTX, complemented by low definition SRT datasets generated from Visium. A detailed characterization of all 37 SRT datasets involved in this study is provided in **Sup. Table 1**, which includes both public data and also data newly generated for this study. Using spacer, we found shared and unique mechanisms that dictate cell infiltration in different tumor types. We characterized intrinsic and extrinsic factors in T cells that impact their localization. To demonstrate its broad usage in another disease context, we studied myocarditis in a mouse model. Overall, spacer made discoveries that would have been overlooked by prior spatial data analysis tools.

### A fully interpretable framework for decrypting spatial cellular localization in tissues

Local cellular neighborhoods within a tumor microenviroment determines whether a T cell is likely to infiltrate. Accurately decrypting the rules for cell infiltration requires a computational framework that is structurally isomorphic to the tissue itself. To this end, we formulated cellular recruitment as a multiple instance learning problem, which mirrors biological reality ^3–5^. We refer to each neighborhood of tumor cells representing the extrinsic microenvironment, defined by drawing a circle in the tumor region based on a pre-set radius, as a “bag” in the language of multiple-instance learning (**Fig. 1ab**). The label of each bag (positive or negative) is defined by the biological ground truth: the presence or absence of an engaging cell (T cell) at its center. The tumor cells within each bag serve as the “instances”, providing the high-dimensional genomic and spatial inputs that drive the recruitment outcome.

Using this conversion of the biological question into a mathematical problem as a starting point, we develop a fully interpretable multiple-instance learning neural network, spacer, to predict recruitment of one type of cells into the tissue microenvironment determined by another type of cells resident in that tissue site. Unlike generic predictors, spacer explicitly models how the transcriptomic landscape of resident cells impacts the infiltration of engaging cells, by investigating the gene expression modality of the resident cells of SRT data. To model the decay of signaling strength over space, we introduce a distance attention module (**Sup. File 1**) to capture signals from the spatial modality of SRT data. Eventually, we build from the bottom up and parameterize an integration module to connect these pieces of information to predict the presence or absence of the engaging cell in the center of the bags, by attending to the transcriptomic modality with attention weights determined by the spatial modality. The structure of the neural network in spacer is fully codified by biological principles and thus the inferred molecular determinants for the cellular interactions are entirely interpretable. Ultimately, spacer outputs the gene-level “spacer scores” that serve as quantifiable readouts of a gene’s attractive or repulsive potential (**Fig. 1c**), effectively identifying the molecular determinants of the cellular makeup of a tissue microenvironment.

**Fig. 1.**
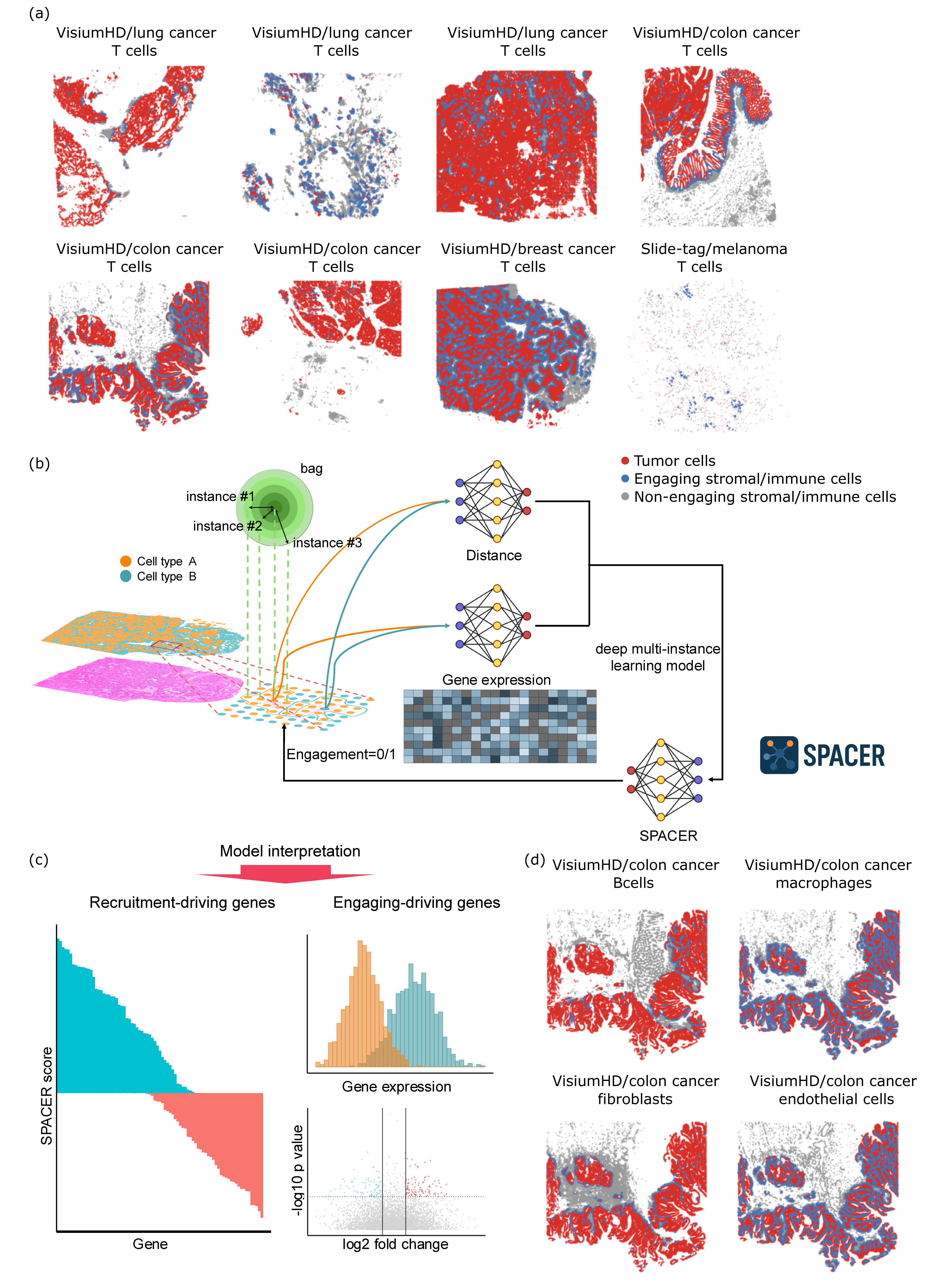
A bottom-up interpretable framework for interpreting rules of cellular localization in tissues. (a) Spatial visualization of the high definition SRT data involved in the human tumor study in the discovery cohort, showcasing the tumor cells and T cells. (b) Cartoon diagram showing raw data, data pre-processing, and the formulation of the spacer model. (c) Biological insights that can be revealed by spacer. (d) Spatial visualization of the colon cancer VisiumHD dataset involved in the human tumor study, showcasing the tumor cells and the other types of stromal/immune cells.

Beyond identifying extrinsic recruitment drivers, spacer enables a systematic dissection of the recruitment-engagement duality. We can apply spacer to predict the label for each defined bag, and we can further compare the true bag label (presence or absence of the engaging cell) with the predicted bag label. Deviation from these true labels would be related to how the intrinsic features of the engaging cells impact their localization, apart from the extrinsic factors from the recruiting cells. We perform differential gene expression and pathway enrichment analyses to uncover these intrinsic features (**Fig. 1c**), which allows us to capture the full spectrum of the recruitment-engagement duality and systematically link both sides of this biological transaction.

Finally, while we used tumor cells and T cells as examples, we are not limited to these cell types. Spacer is widely applicable for the other infiltrating stromal and immune cells as well in the tumors (**Fig. 1d**), and also in other contexts where the migration of cells is based on environmental cues.

### Determinants of T cell recruitment in human tumors

We applied spacer to 7 VisiumHD and 1 Slide-Tag, as well as 20 Visium datasets in our tumor discovery cohort, which span several tumor types (**Fig. 1a**). As spacer was applied to these datasets collectively, we expected spacer to extract general pan-cancer mechanisms. To interpret what spacer learned, we first examined the genes expressed in the tumor cells that were inferred to recruit T cells (**Fig. 2a** and **Sup. Table 2**). Strikingly, we found that several of the top genes are related to the HLA machinery or are known to be well-known tumor antigens (**Fig. 2a**). Class I HLA-related genes, including *HLA-A*, *HLA-B*, *HLA-C*, and *TAP2*, appeared to be more dominant in the genes with top spacer scores (explained above), as opposed to class II genes. We next employed GOrilla ^6,7^ to identify the enriched pathways in these genes in an unbiased manner (**Fig. 2b**), which showed that many immune response-related terms and antigen presentation-related terms, especially for class I antigen presentation, were significantly enriched.

**Fig. 2.**
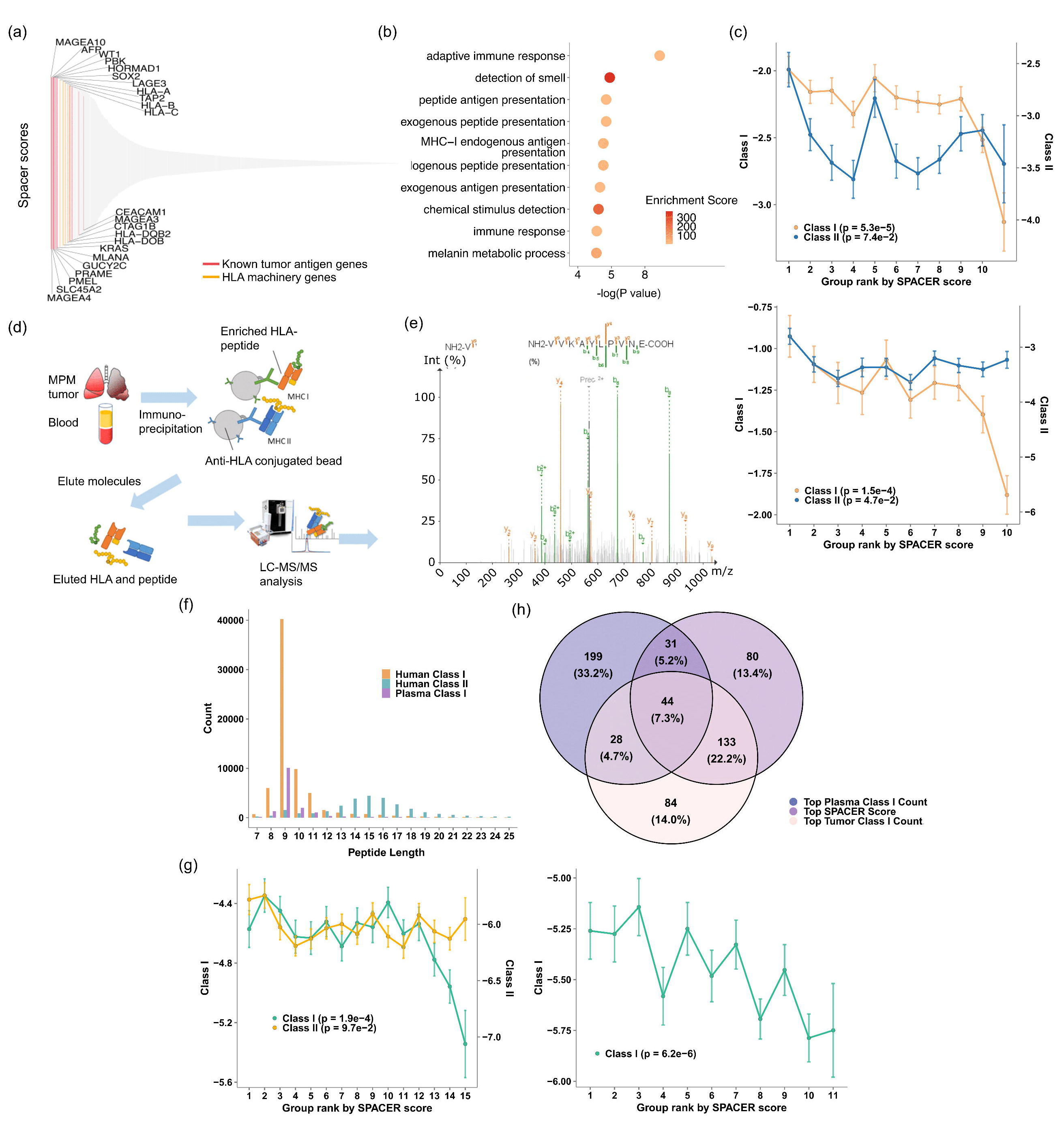
Determinants of T cell recruitment in human tumors. (a) Spacer results of the tumor cell-expressed genes that positively promote T cell recruitment. (b) GO analysis of the ranked genes from (a). (c) The top genes predicted to promote T cell recruitment also encode more immunogenic peptides. SysteMHC (left) and IEDB (right). The ranked genes from (a) were evenly divided into 10 bins. The Y axis shows log((peptide counts+1)/protein length) for each gene. (d) Schematic overview of the immunopeptidomics pipeline. (e) Representative tandem mass spectrum (MS/MS) of an immunopeptidome bound to MHC-I. The peptide VIVKAYLPIVNE (from protein EF2) was identified and sequenced based on its fragmentation pattern using tandem mass spectrometry. The spectrum shows b-ions (green) and y-ions (orange) resulting from peptide backbone cleavage. Fragment ion peaks are annotated accordingly, with the precursor ion (2+) also indicated. The sequence shown at the top represents the N-terminal to C-terminal orientation (left to right), and the matching b- and y-ion series confirm high-confidence peptide identification. (f) Peptide length distributions of the class I and II immunogenic peptides identified from the tumors and plasma. (g) The top genes predicted to promote T cell recruitment encode more immunogenic peptides. Human tumors (left) and plasma (right). (h) Overlap of top genes ranked by spacer scores and immunopeptidomics datasets. Genes were ranked by spacer scores, tumor MHC class I peptide counts, and plasma MHC class I peptide counts. Top 25% of genes from each ranked list were selected for drawing the Venn diagram.

Tumor antigens encode immunogenic peptides presented by MHCs that induce T cell reactivities. To determine if top spacer score genes encode more immunogenic peptides, we quantified the number of peptides encoded by each of these genes’ protein products that are also known to be binding to at least one HLA, according to immunopeptidomics data from IEDB ^8^ or SysteMHC ^9^. Since this SRT data came from a mix of genetic backgrounds without HLA information, we took a HLA-agnostic approach to match immunogenic peptides. We normalized the numbers of matched peptides by each gene by the length of the full length proteins. The genes with higher spacer scores were also likely to encode more class I immunogenic peptides normalized by protein length, while the trend is weaker for class II immunogenic peptides (**Fig. 2c, left** (SysteMHC) and **Fig. 2c, right** (IEDB)). For further validation, we also generated immuno-peptidomics data from 40 pleural mesothelioma (PM) patients (**Fig. 2de**). Class I and II immuno-peptidomics data were generated from the tumor sites, and class I immuno-peptidomics data was generated from plasma. As expected, most class I immunogenic peptides were 8-11 amino acids in length, while class II immunogenic peptides were 15 amino acids in length on average (**Fig. 2f**). Matching the peptides encoded by each gene to the PM tumor immuno-peptidomics data also revealed that top spacer score genes encode more class I immunogenic peptides, while the trend is less obvious for class II (**Fig. 2g, left**). The same trend was also observed by matching to the immunogenic peptides found in the patient plasma (**Fig. 2g, right**). This was expected as the tumor and plasma immunopeptidomics data are both searched against the same reference genome, therefore cells in the tumors and the plasma shall encode the same proteins and same peptides that are presentable by the same MHCs for the most part. However, we should also likely observe higher concordance between the tumor immunopeptidomics data with spacer results, as opposed to the blood immunopeptidomics data, due to tumor-specific presentation of peptides on MHCs. Indeed, the overlap of peptides from genes of the top 25% spacer scores with immunogenic peptides from the tumor sites is much higher than the overlap with immunogenic peptides from plasma (**Fig. 2h**). The hyper-geometric test P val. for the overlap between spacer top genes and tumor immunogenic genes was 3.3E-10, while >0.05 for the other two overlaps, confirming that spacer is capturing putative antigen genes that are tumor specific. In summary, while HLA-presented immunogenic epitopes are known to be T cell recognition targets, we showed for the first time that their presence in local tumor niches also impacts the recruitment and spatial patterns of T cell distributions. This had not been previously demonstrated by existing SRT data analysis tools.

### Tumor factors that inhibit T cell recruitment

Similarly, we sought to determine the tumor-expressed genes that negatively impact the recruitment of T cells into tumors. Many of the top genes are associated with extracellular matrix (ECM) components (**Fig. 3a** and **Sup. Table 2**). In fact, the gene with the highest spacer score is *MUC5B* (**Fig. 3a**), which encodes a secreted gel-forming mucin. GOrilla analyses also confirmed that ECM terms such as collagen metabolic processes and O-glycan-related factors were significantly enriched (**Sup. Fig. 1**). In particular, O-linked glycans are built by Golgi apparatus in mucins, which are then packaged into secretory granules ^10^. We also examined the cellular location of the proteins encoded by human genes (definition in method section, and also see **Sup. Table 3**). The top genes that were inferred to repel T cell infiltrations encoded more proteins that are extracellularly located - either plasma membrane-associated or secretedin comparison with genes that were inferred to positively attract T cell infiltrations (**Fig. 3b**). These observations suggest that the characteristics of the ECM surrounding tumor cells might influence T cell infiltration. We then explicitly examined the expression of ECM proteins (**Fig. 3c**). The expression of genes encoding ECM component proteins were more highly expressed in the tumor regions that are less infiltrated by T cells, compared with the non-infiltrated regions. The largest difference occurs for mucins (Wilcox test P val.<0.001). Interestingly, while these ECM components can be secreted by many types of cells found in the tumor microenvironment (tumor cells, cancer associated fibroblasts, *etc*), mucins are known to be mainly secreted by tumor cells ^11,12^. The higher expression of mucins in tumor cells vs. stromal/immune cells was also confirmed in our own SRT data (**Fig. 3d**). In contrast, the expression of collagens showed the opposite trend (Wilcox test P val.<0.001).

**Fig. 3.**
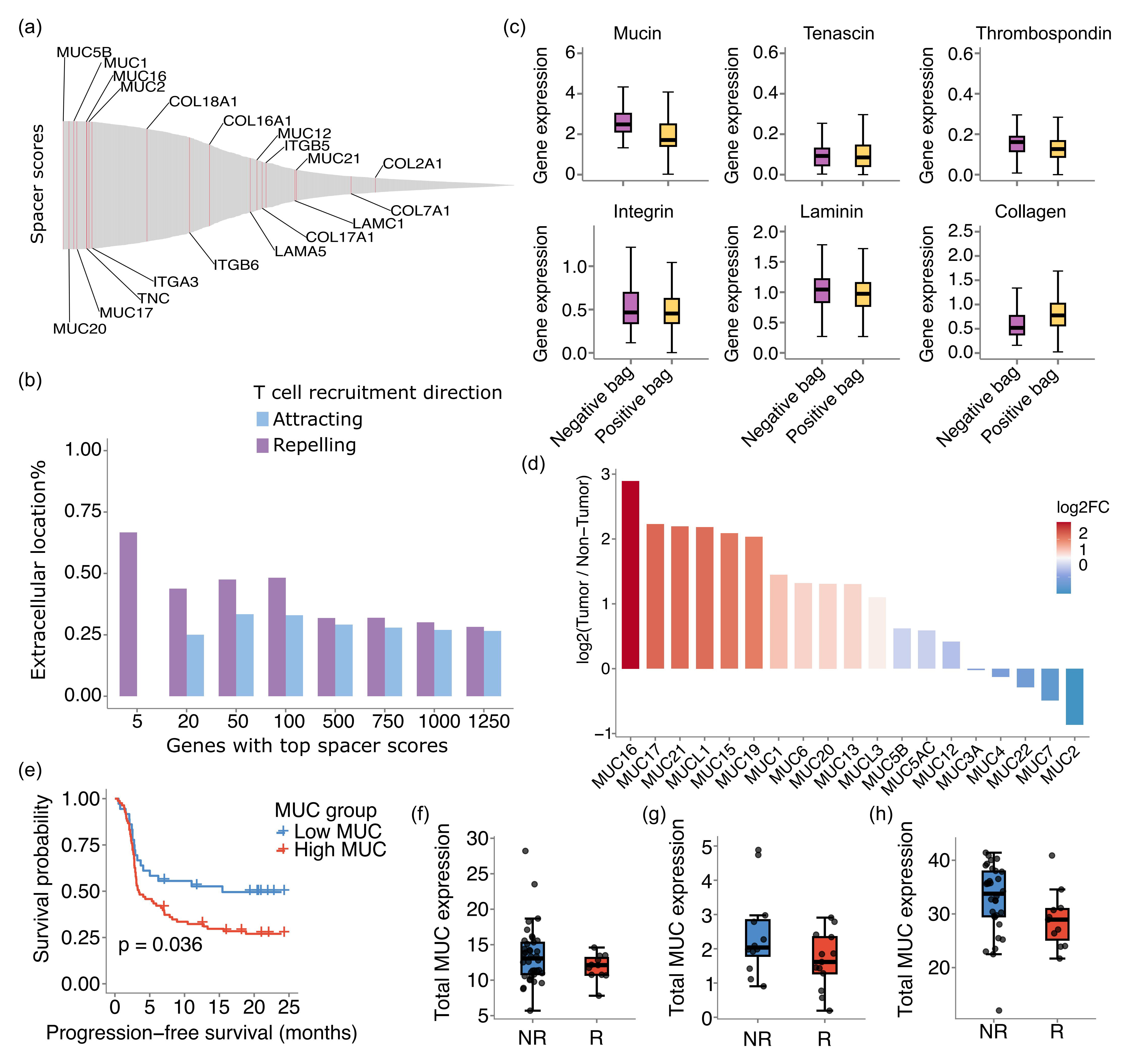
Factors determining inhibition of T cell recruitment. (a) Spacer results of the tumor cell-expressed genes that repel T cell recruitment. (b) The top tumor cell-expressed genes that are predicted by spacer to repel T cell recruitment encode more extracellular proteins than intracellular proteins. The top tumor cell-expressed genes that are predicted by spacer to attract T cell recruitment serve as a control. (c) The expression of extracellular matrix genes in the positive and negative bags (neighborhoods of tumor cells with and without T cell infiltrations) of the spacer analyses. (d) The higher expression of mucins in tumor cells over stromal/immune cells is also confirmed in our own SRT data. (e) Kaplan-Meier estimator for PFS of the Liu *et al* cohort, where the ICI-treated patients were dichotomized based on mucin gene expression. (f-h) Mucin gene expression in the ICI-treated responder and non-responder patient subsets of the Hugo *et al* (f, unit=log(TPM)), Riaz *et al* (g, unit=log(FPKM)), and Kim *et al* (h, unit=log(TPM)) cohorts. R=responder; NR=non-responder.

Given these observations, we asked if mucin overexpression is associated withimmune-checkpoint inhibitor (ICI) resistance. We first evaluated the Liu *et al* ^13^ cohort where there are a total of 119 melanoma patients on anti-PD1 treatment and also with pre-treatment RNA-sequencing data. As is shown in **Fig. 3e**, over-expression of mucins is indeed associated with worse progression free survival (PFS) according to our analyses (P val.=0.02). Next, we evaluated the Hugo *et al* ^14^ cohort (anti-PD1, 25 melanoma patients with complete data), the Riaz *et al* ^15^ cohort (anti-PD1/anti-CTLA4, 49 melanoma patients with complete data), and the Kim *et al* ^16^ cohort (anti-PD1, 43 gastric cancer patients with complete data). Time-to-event data were not available for these cohorts, so we evaluated the expression levels of mucins in the responders and non-responders. As is shown in **Fig. 3f-h**, the non-responders have higher mucin expression compared to responders (P val.=0.038 for the Hugo *et al* cohort in **Fig. 3f**; P val.=0.046 for the Riaz *et al* cohort in **Fig. 3g**; P val.=0.41 for the Kim *et al* cohort in **Fig. 3h**).

### Shared and unique cellular recruitment mechanisms across tumor and tissue types

Using spacer, we asked if there are differences between immune cell recruitment mechanisms between tumor types. In **Fig. 4a**, we showed the expression of tumor genes in all our 10 high and 20 low definition tumor VisiumHD/Slide-Tag datasets (**Sup. Table 1**, including two additional VisiumHD 3’ datasets), ranked by the spacer scores from **Fig. 3**. The genes on the top were predicted to be the most significant drivers of T cell recruitment. First, data sets of the same cancer type tended to cluster together, especially for melanoma, breast, prostate and colon cancers. This suggested there might be T cell recruitment mechanisms that are unique and conserved to each cancer type. Interestingly, many instances of SRT datasets of the same cancer type but different resolution levels clustered together, validating our approach of merging these data. Then we examined tumor expressed genes, ranked by spacer, and found that the very top ranking genes (labeled in red on the Y axis of **Fig. 4a**) have low average expression across tumor cells, almost uniformly across all datasets. Then the next segment of genes (labeled in green) with weaker impact on recruitment of T cells generally have higher gene expression in the tumor cells. The genes (orange) at the bottom have moderate gene expression. As spacer was applied to all datasets of all cancer types together, the genes that rank on the very top (red genes) are likely to be pan-cancer drivers of T cell infiltration with the most potent effect. And our results suggest that this core set of genes tend to be down-regulated by tumor cells in general, most likely as a mechanism of avoiding immunosurveillance.

**Fig. 4.**
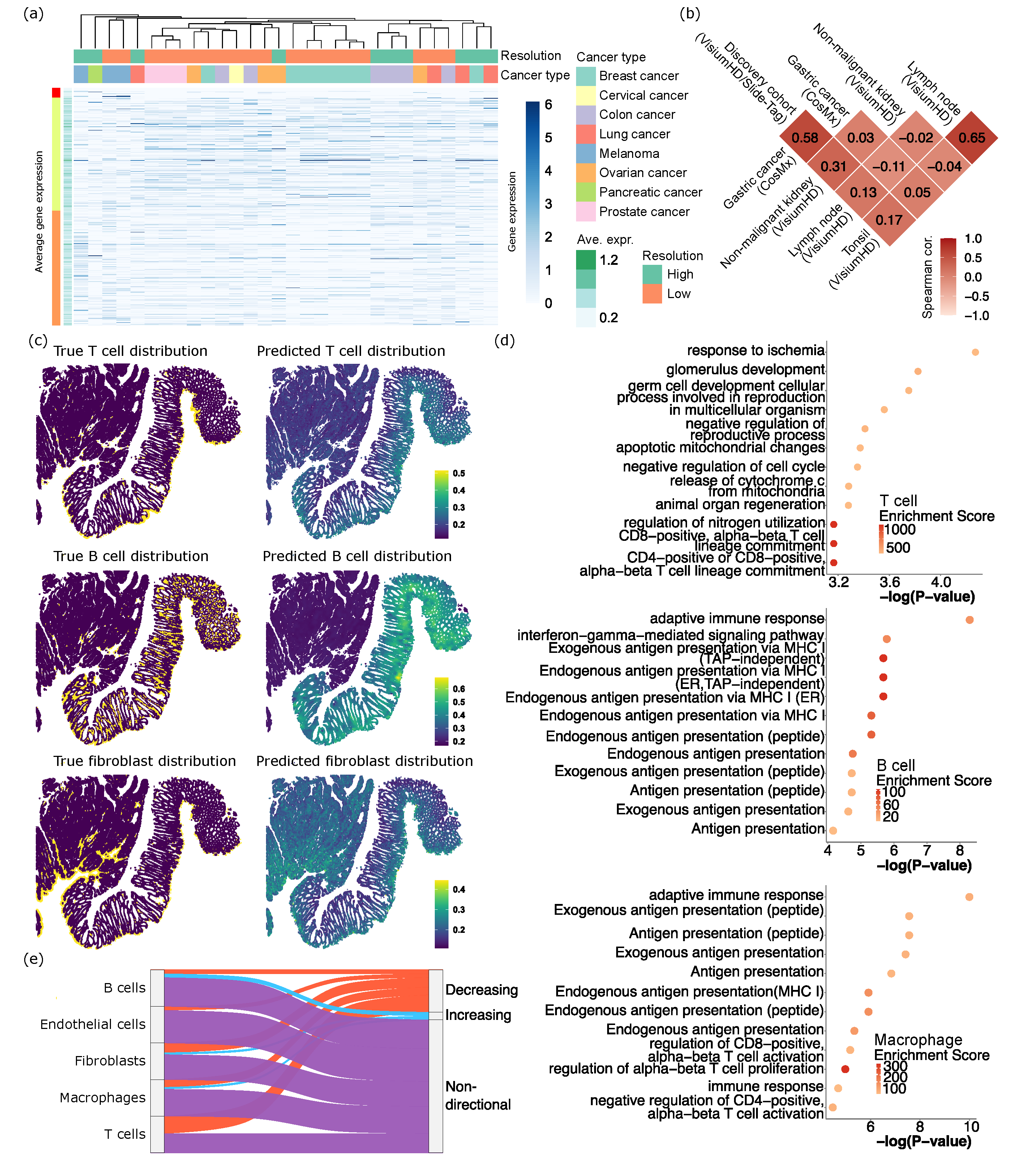
Shared and unique recruitment mechanisms across tumor types and engaging cell types. (a) Expression of tumor cell genes that promote T cell infiltrations, in each of the high and low resolution SRT datasets involved in our study. The main heatmap shows the expression in each dataset, while the bar towards the left shows averaged expression across datasets. The columns (datasets) were ordered through hierarchical clustering based on tumor cell gene expression, and labeled by the resolutions and cancer types of the datasets. (b) Spearman correlations between the spacer scores that we computed from the human tumor discovery cohort (VisiumHD/Slide-Tag, n=8), Gastric cancer cohort (CosMx, n=2), non-malignant kidney sample (VisiumHD, n=1), lymph node sample (VisiumHD, n=1), and tonsil sample (VisiumHD, n=1). (c) The true and spacer-predicted T cell, B cell and fibroblast distribution in one ovarian cancer VisiumHD dataset. (d) GO analyses of the top tumor cell-expressed genes predicted by spacer to promote stromal/immune cell infiltrations. Also refer to **Sup. Fig. 2cd**. To emphasize the differences between the different types of cells, for the list of ranked genes of each cell type for its GO analysis, we removed genes that also ranked as high in the ranked gene lists of any other cell types. (e) The percentages of the top spacer score genes for each type of stromal/immune cells that show increasing, decreasing, and non-directional expressional changes over developmental stages. Data from TEDD 2.0 ^18^.

To further evaluate how much the T cell recruitment mechanism is shared or is unique across different tumors and samples, we generated 2 CosMx datasets from gastric cancers. As CosMx is imaging-based, while VisiumHD, Slide-Tag and VisiumHD 3’ are all sequencing-based, we decided not to analyze the CosMx data using the fine-tuning approach as above, due to compatibility issues. Instead, we directly applied spacer to these samples, and then calculated the correlation between the spacer scores from the human tumor discovery cohort and the CosMx cohort (**Fig. 4b**). The Spearman correlation reached 0.58, which again suggests the existence of a pan-cancer program in T cell recruitment, while the inter-tumor differences are also substantial. Also, examining the T cell-recruiting genes detected by spacer (**Sup. Table 2**), we found that the MHC machinery genes (*HLA-A*, *-B*, *-C*, *-DRB3*, *-DOB*, *-DQB2*) and type I interferon signaling pathway genes (*e.g. IFI6*) ranked even higher compared to our human tumor discovery cohort. This result, combined with our observations above, indicates that, even for cancer types that are generally less immunogenic (*e.g.* gastric cancers), T cell infiltration is still influenced by antigen presentation in the local tumor microenvironment.

On the other hand, we applied spacer to three more VisiumHD datasets generated from non-malignant kidney, lymph node, and tonsil (**Fig. 4b**). In the kidney, T cells infiltrated into the tubular areas surrounding the glomeruli, suggesting inflammation in the kidney epithelium in this biospecimen (**Sup. Fig. 2ab**). Therefore, we applied spacer to study the infiltration of T cells into the kidney epithelium. Lymph nodes and tonsils are secondary lymphoid organs so T cells are resident cells, as opposed to “infiltrating” these organs. Thus, we deployed spacer here as a negative control. And as expected, while the correlation between lymph node and tonsil was high, the correlation between lymph node/tonsil and human tumors/kidney was close to 0 (**Fig. 4b**). There is a weak positive correlation between human tumors and kidney, indicating that inflamed non-malignant tissues and tumors share some similarities in the biological mechanisms that determine the spatial organization of infiltrating T cells.

### Comparing across different engaging cell types

We next focused on interactions between different cell types. In ovarian cancer, we showcased the true and also spacer-predicted spatial distributions of T cells, B cells, and fibroblasts (**Fig. 4c**). These three cell types demonstrate differential patterns of tumor infiltration, suggesting that stromal and immune cell infiltrations are diverse. The spacer-predicted distributions also closely follow the true spatial distributions, indicating that spacer has successfully captured these differential mechanisms. We next used GOrilla to analyze the significantly enriched pathways in the tumor-expressed genes that were inferred to drive stromal and immune cell infiltrations. We focused on five cell types, including T cells, B cells, macrophages, fibroblasts and endothelial cells. For T cells, we observed enrichment of development and wound healing terms (**Fig. 4d**), in addition to GO terms related to T cell activation. For B cells and macrophages (**Fig. 4d**), we observed enrichment of antigen presentation-related GO terms. For fibroblasts (**Sup. Fig. 2c**), we observed an enrichment for synaptic function terms. For endothelial cells, we observed melanocyte development-related terms (**Sup. Fig. 2d**). Melanoma is highly vascularized and other vascularized cancer types could also demonstrate similar expression patterns, which might be conducive to endothelial cell engagement and angiogenesis. Nevertheless, GO analyses show that the signals for the recruitment of T cells, B cells and macrophages are overall much stronger, compared with fibroblasts and endothelial cells (compare P val. of **Fig. 4d** with **Sup. Fig. 2cd**; further discussion in **Sup. File 2**).

We observed differential GO enrichment between T cells and B cells/macrophages. The enrichment of GO terms related to antigen presentation for B cells and macrophages could be related to the antigen presentation functions of B cells and macrophages themselves ^17^. On the other hand, the enrichment of developmental and wound healing terms for T cells is intriguing. We conjecture that this could be related to the fact that many tumor antigen genes are expressed only early on in a human being’s developmental processes, so T cells would mount a response once they see these genes re-expressed again in tumors. To prove this hypothesis, we examined the top genes that spacer inferred to be positively impacting the recruitment of each type of stromal/immune cell types (**Fig. 4e**). We validated these genes with the temporal gene expression data from Temporal Expression during Development Database ^18^, with which we calculated the average expression of each gene in fetal, childhood and adult stages. We defined genes as “decreasing” if their expression is highest in fetal stages and lowest in adult stages, and *vice versa* as “increasing”. Genes with no consistent trend are labeled as “non-directional”. As **Fig. 4e** shows, the top genes with the strongest T cell recruitment potential indeed possess the largest proportions of “decreasing” genes, consistent with our hypothesis. These “decreasing” genes include well known tumor antigen genes like *PMEL* ^19^ and *AFP* ^20^, but also include genes that are less well known, such as *PBK*, which might be a new target for T cell-based immunotherapy.

### Advantages of SRT in yielding insights on cellular infiltration into tissues

To our knowledge, there have been no other tools developed for inference and interpretation of cellular recruitment and localization from SRT data. Therefore, we cannot execute a benchmark study comparing spacer with other software. However, to empirically demonstrate the advantage of considering the exact locations of the infiltrating cells provided by SRT, we analyzed the bulk RNA-sequencing data from the The Cancer Genome Atlas Program (TCGA) project, for all cancer types combined (**Sup. Fig. 3**). We examined the top 50 spacer score genes, and calculated the correlation between these genes’ expression from the bulk sequencing data with the predicted infiltration of CD4^+^ (expression of *CD4*) and CD8^+^ T cells (sum of expression of *CD8A* and *CD8B*). As **Sup. Fig. 3** shows, we observed nearly no positive correlation overall when we aggregate over all cancer types and all genes. This result attested to the advantages of SRT in yielding insights on cellular infiltration into tissues. Spacer analyzes whether T cells actively localize according to tumor cell gene expression in local niches, which should yield more accurate insight into T cell responsiveness, benefiting from the higher granularity of the SRT data.

### Spacer elucidates the mechanisms of T cell engagement with tumor cells

We studied the mechanisms of T cell engagement with the tumor cells, conditioned upon the spacer-predicted recruitment patterns of the infiltrating T cells. Above, spacer was essentially trained to predict T cell infiltration based on nearby tumor cell gene expression. Then we used this trained spacer to infer the tumor cell regions where T cells should and should not infiltrate. We examined the differential gene expression of the T cells found in the “predicted to infiltrate” regions (therefore a correct prediction) and “predicted not to infiltrate” regions (an incorrect prediction), with the genes and their expression shown in **Sup. Table 4**. We hypothesize that this analysis would give us insight into both the transcriptomic changes of T cells upon engagement with the tumor cells, and also the reasons of why some T cells still infiltrated regions where there is a lack of tumor-derived recruiting signals (such as lack of expression of tumor antigens and HLA genes).

We first performed a GO analysis to identify the enriched pathways in the genes that are differentially regulated (**Fig. 5a**). As expected, we identified pathways that are related to immunity and T cell activation, and also pathways that are related to migration of T cells. We then examined one specific VisiumHD dataset from colon cancer, and performed network analyses of the T cells’ gene expression using hdWCGNA ^21^. This analysis revealed several modules of genes with highly correlated expression in each module (**Fig. 5b**). The *MAPK* signaling module, which is one of the core pathways for T cell activation ^22,23^, and the T cell migration module contained genes that are among the top genes with the largest expressional differences between the “predicted to infiltrate” and “predicted not to infiltrate” regions in this dataset (**Fig. 5c**).

**Fig. 5.**
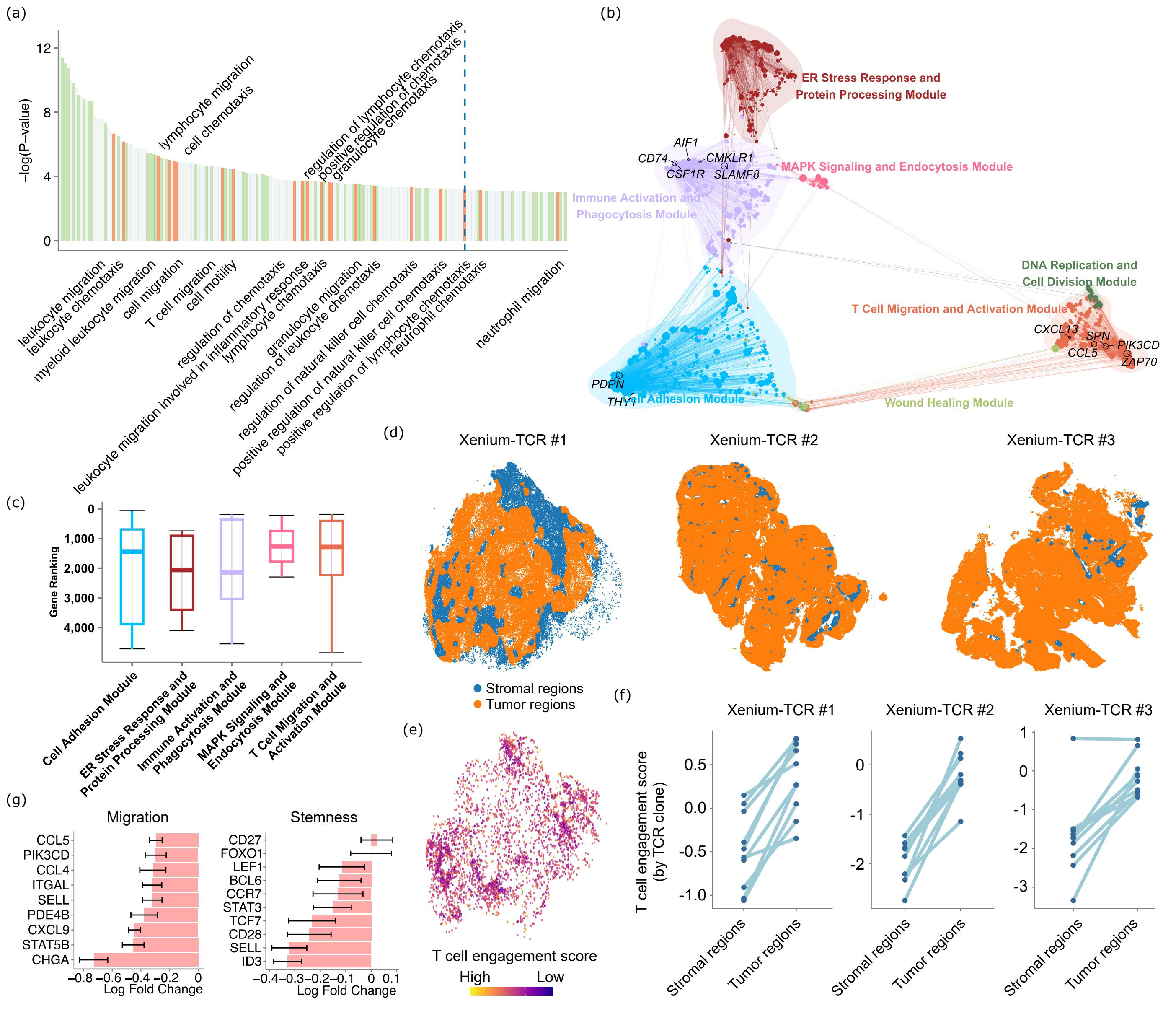
Spacer interprets the mechanisms of T cell engagement with tumor cells. (a) GO analyses of the genes that are differentially expressed between the “predicted to infiltrate” tumor regions and “predicted not to infiltrate” regions. (b) Gene networks detected in the differentially expressed genes as in (a) but for the colon cancer VisiumHD dataset. (c) The ranks of the genes in each gene module (b) that entered the differential expression analysis for the colon cancer VisiumHD dataset, with the genes ranked by the absolute values of the log fold changes. (d) Visualization of the three Xenium-TCR-seq datasets, with the stromal and tumor regions colored differently. (e) The visualization of the spacer-predicted tumor-specific T cell engagement score in the spatial context of the first Xenium-TCR-seq dataset. (f) The T cell engagement scores of the T cells from the top TCR clonotypes of each Xenium-TCR-seq dataset, in the respective stromal and tumor regions. (g) Differential gene expression analyses of T cell migration- and stemness-related genes between the “predicted to infiltrate” regions and the “predicted not to infiltrate” regions. Negative values indicated higher expression in the “predicted not to infiltrate” regions.

We hypothesize that these differentially expressed genes in the T cells could reflect the transcriptomic changes of T cells upon engagement with the target tumor cells, namely indicating specificity of T cells against tumor cells. For example, we observed that many type I Interferon response genes are among the top genes that are most highly expressed in the “predicted to infiltrate” regions as opposed to the “predicted not to infiltrate” regions, including *IOAS3*, *MX1*, *IFI27*, *ISG15*, *IFITM3*, *XAF1*, *OAS1*, *IRF7*, *OAS2* (**Sup. Table 4**). We recently developed a Xenium-TCR-sequencing technology (preprint ^24^), where TCR-specific probes were added to the Xenium panel so that expression of close to 500 genes and the presence of about 200 TCR clonotypes can be captured concurrently for each dataset. We sought to validate that the signature indeed indicates tumor-reactiveness, by investigating three datasets that we generated *via* Xenium-TCR-seq. For these datasets, we visualized the cell typing results as well as the presence of TCR clones in the spatial context in **Sup. Fig. 4**. We divided the whole slide sections into the tumor regions and the stromal regions, based on the concentrations of the tumor cells and stromal/immune cells in each physical location (**Fig. 5d**). Then we calculated a tumor-reactive T cell signature for all the T cells found in these three datasets by taking the sum of the products of the expression of each T cell gene captured by Xenium-TCR and the log fold change of the same gene from the differential expression analyses of “predicted to infiltrate” *vs.* “predicted not to infiltrate” above. We visualized the signature scores for the T cells of one of our Xenium-TCR-seq datasets in their spatial context in **Fig. 5e**. For ease of visualization and analysis, we further focused on the top most expanded TCR clones of each dataset and compared the average signature scores of the T cells of each clonotype in the tumor fraction and in the stromal fraction. As we suspected, the signature scores are significantly higher in the tumor fractions compared with the stromal fractions (T test p val.=9E-6, 2.3E-5, and 1.9E-4, respectively) (**Fig. 5f**), confirming the tumor-reactive gene signature. Importantly, the analysis introduced here is not merely a clearer visual presentation compared with **Fig. 5e**. Rather, the different T cells found in the same TCR clonotype share the same TCR. By controlling for the factor of TCR clonotypes (so the T cells should have the same reactivities towards the exposed antigens), we were able to exclude the extrinsic factors and focus on the intrinsic factors of the T cells that impacted tumor-reactivity.

While the genes that are highly expressed in the T cells in the “predicted to infiltrate” regions are related to immune and defense responses, this does not yet completely explain the enrichment of the migration signatures in the GO analyses (**Fig. 5a**). In particular, the differential gene expression analyses of **Fig. 5ab** are agnostic of the direction of gene expression. Therefore, we hypothesize that the T cells that are infiltrating the “predicted not to infiltrate” regions are in fact the T cells with high migration potentials. To test this hypothesis, we examined a number of marker genes known to be associated with higher T cell migration capabilities (**Fig. 5g**), and they are all indeed more highly expressed in the T cells in the “predicted not to infiltrate” regions, compared with the T cells in the “predicted to infiltrate” regions. Therefore, T cells observed in the tumor regions, where the tumor cells’ expression does not support T cell recruitment, are more migratory inherently, rather than responding to tumor cells specifically. Ancillary to this result, we also examined a set of marker genes for T cell stemness (**Fig. 5g**). These genes are also more highly expressed in the “predicted not to infiltrate” regions overall. Stem-like T cells are known to continuously seed into tumors from tumor-draining lymph nodes and drive more sustained anti-tumor responses than terminally differentiated/exhausted T cells ^25,26^. It has also been reported that stem-like T cells tend to be sequestered away from antigens ^27,28^, which echoes our observation.

Importantly, spacer revealed the “outcome” of anti-tumor response for T cells in the “predicted to infiltrate” regions but the “cause” of T cell presence in the “predicted not to infiltrate” regions, which advances beyond prior T cell tumor-reactive signature works ^29^

### Spacer reveals unexpected dominance of CD4^+^ T cells in heart myocarditis

To demonstrate the universal applicability of spacer beyond oncology, we next deployed spacer to a different biological system in a different species, to demonstrate its broad usage. We generated one male and one female *Pdcd1*^−/−^*Ctla4*^+/–^ mice, which can recapitulate clinicopathological features of severe immune checkpoint inhibitor (ICI)-induced myocarditis (ICI-MC) ^30^. We generated VisiumHD data from these two mice. **Fig. 6a** shows the H&E staining images, the typed cells in their spatial context, and also the typed cells in their UMAP space. Then we focused on the recruitment of T cells into the ventricular cardiomyocytes, as these cells are known to be the primary targets of the immune system during myocarditis ^31–33^.

**Fig. 6.**
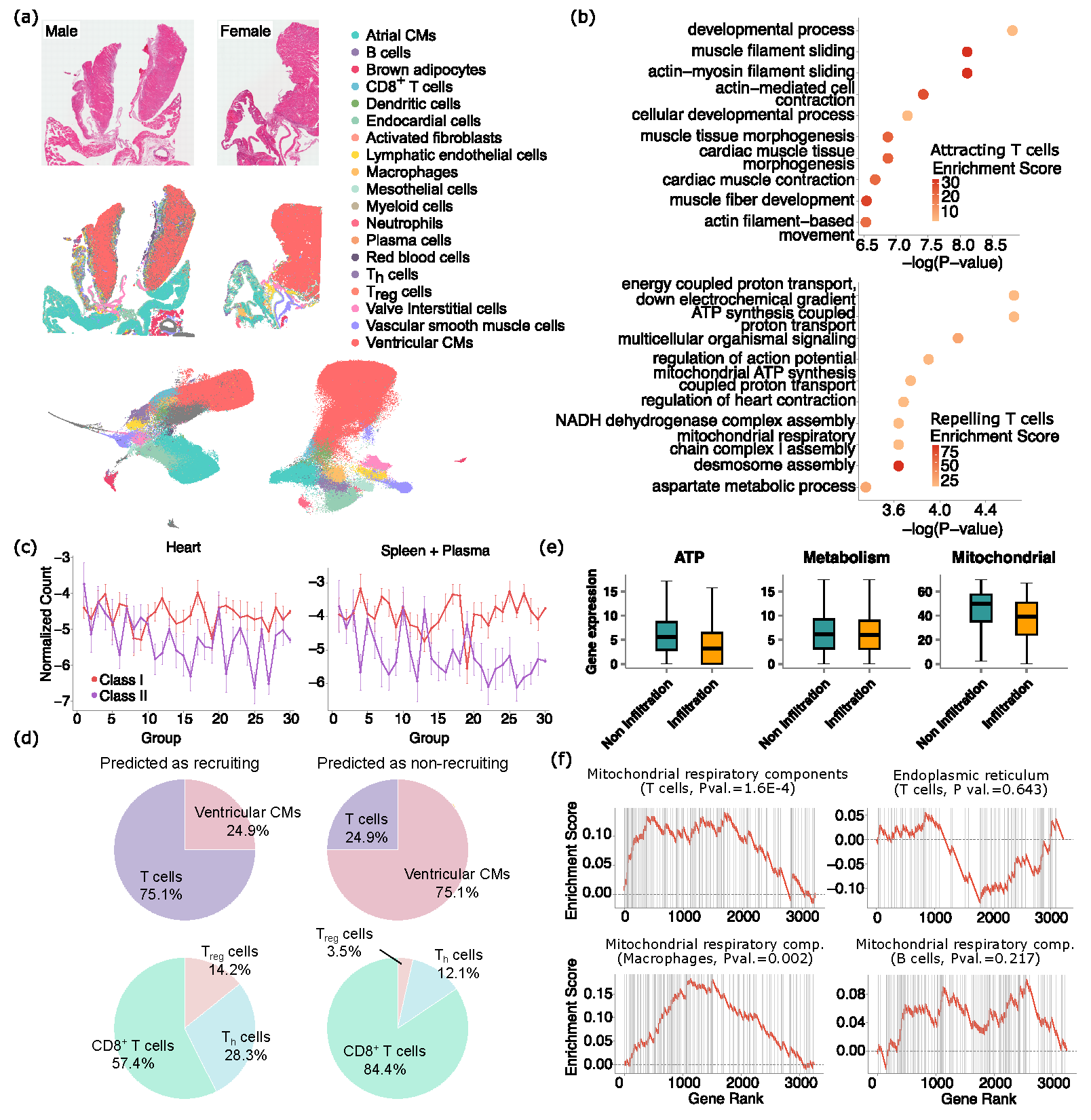
Spacer reveals involvement of CD4^+^ T cells in heart myocarditis. (a) The H&E staining images and cell type labels of the cells in their spatial context, of the mouse myocarditis VisiumHD datasets, as well as the UMAP plots of the cells and cell type labels. (b) GO analyses of the ranked cardiamyocyte genes that are predicted by spacer to promote or inhibit T cell engagement in the ventricular cardiomyocyte regions. (c) The top genes predicted to promote T cell recruitment encode more class II immunogenic peptides, while the trend is much less clear for class I immunogenic peptides. Immunopeptomics data collected from heart (left) and spleen/plasma (right). (d) The proportions of cardiomyocytes and T cells (top) and the proportions of the different types of T cells out of total T cells (bottom), in the ventricular cardiomyocyte regions that are predicted by spacer to be T cell infiltrated or non-T cell infiltrated. (e) The gene expression levels of ATP-related genes, metabolism-related genes, and mitochondria-related genes in the “predicted to infiltrate” and “predicted not to infiltrate” regions. (f) T cell genes that are more highly expressed in the “predicted not to infiltrate” regions than the “predicted to infiltrate” encode proteins that are enriched in the mitochondrial respiratory components. The enrichment of endoplasmic reticulum and expression of genes in macrophages and B cells serve as negative controls. The TCA cycle-related mitochondria components include the mitochondrial matrix, the integral component of mitochondrial inner membrane, the mitochondrial intermembrane space, and the inner mitochondrial membrane protein complex. Annotations of cellular compartments are from the COMPARTMENTS database ^43^.

We first interpreted the genes that were inferred by spacer to be positively correlated with the infiltration of T cells (**Sup. Table 2**). We observed *Myh6* among the top genes, which was known to encode MHC-presented peptides that induce T cell infiltrations in heart myocarditis ^30^. We observed *Tnni3*, which is a known marker of cardiomyocyte damage during myocarditis ^34^. *H2-Q7*, *H2-K1*, *H2-Ab1*, and *H2-Q6* are also ranked among the top, which encompass both class I and II MHCs. We did not identify any tumor antigen genes, as was the case for the application of spacer in human tumors above, which indicates the specificity of spacer in detections of cellular localization signals. However, GOrilla analyses (**Fig. 6b**) revealed that developmental and morphogenesis processes are enriched in the cardiomyocyte-expressed genes that promote T cell infiltration. These results echo **Fig. 4de**, where the tumor genes that promote T cell infiltration are found to be enriched in developmental pathways. Related to this, *Myh6* is known to be abundantly expressed in the cardiac ventricles during embryonic development, while, following birth, cardiac ventricles predominantly switch to express *Myh7* ^35^. Motivated by these results and mirroring our tumor study, we further performed immunopeptidomics experiments on the mouse heart tissues, as well as in the blood and spleen tissues. As **Fig. 6c** shows, we curiously found that the top cardiomyocyte genes identified by spacer to recruit T cell infiltration tend to encode more class II immunogenic peptides with significance achieved (P val.=0.019) and barely more class I immunogenic peptides (P val.=0.067), with the immunogenic peptides profiled from the heart. For immunogenic peptides profiled from the spleen and blood combined, we observed a similar trend with weaker statistical significance (P val.=0.076 for class II and P val.=0.243 for class I). These results suggest that there are likely only a few immunogenic MHC I epitopes that dominate Cd8^+^ T cell response, consistent with prior observations ^36^, while there are probably many more MHC class II epitopes that induce Cd4^+^ T cell responses, which is reflected by the significant enrichment captured in **Fig. 6c**.

While Axelrod *et al* ^30^ has reported critical roles that Cd8^+^ T cells play in myocarditis and we also observe abundant Cd8^+^ T cell infiltration in ventricular cardiomyocytes (**Fig. 6d**), our results raise the curious possibility that the MHC class II-Cd4^+^ T cell axis might also be active in myocarditis. We examined the ventricular cardiomyocyte regions in the two mice and applied spacer to predict where T cell infiltrations should or should not occur (**Fig. 6d**). The predicted “should infiltrate” regions indeed have proportionally more T cells than the “should not infiltrate” regions, which is expected. However, when we examined the different subtypes of T cells, including Cd8^+^, T_h_, and T_reg_ cells, out of total T cells, we found that Cd8^+^ T cells actually decreased in proportions in the “should infiltrate” regions, while T_h_ and T_reg_ cells selectively increased. This result provides compelling evidence to support the hypothesis that the MHC class II-Cd4^+^ T cell axis is functional and could even be more active than the MHC class I-Cd8^+^ T cell axis during myocarditis.

To interpret the genes that were inferred by spacer to be negatively correlated with T cell infiltration, we performed GO enrichment analysis and discovered ATP- and TCA cycle-related terms (**Fig. 6b**). We further examined the gene expression in the cardiomyocyte regions with or without T cell infiltrations. We found that ATP- and mitochondria-related gene sets are indeed more highly expressed in the regions without T cells, but this is not true for the expression of general metabolism-related genes (**Fig. 6e**, Wilcox test P val. <1E-3, =0.018, and <1E-3 from left to right). These results suggest that the correlation is specific to the mitochondria where the TCA cycle happens. We re-affirmed this observation by also performing Gene Set Enrichment Analysis (GSEA) ^37^, to test the enrichment of the expression of the top spacer score genes in the components of mitochondria involved in the TCA cycle, other components of the mitochondria, and also the endoplasmic reticulum as a control (**Fig. 6f** and **Sup. Fig. 5**). We also performed the same spacer analyses for macrophages and B cells, and observed similar results for macrophages with statistical significance achieved, but not for B cells (**Fig. 6f** and **Sup. Fig. 5**). These results suggest that T cells and macrophages are the major effectors of tissue damage in myocarditis.

### Functionally active CD4^+^ T cell responses in heart myocarditis

We further pursued the possibility that the MHC class II-Cd4^+^ T cell axis could be functionally active, despite their smaller proportions out of total T cells. We first examined the activation, exhaustion and memory expression signatures of the Cd8^+^, T_h_, and T_reg_ cells in the SRT data, in the predicted “should infiltrate” regions, compared with the “should not infiltrate” regions of the cardiac ventricles. The Cd8^+^ T cells are less activated and more exhausted (**Fig. 7a**). On the other hand, the T_reg_ cells showed a trend of being more activated and more exhausted in the “should infiltrate” regions, without statistical significance (**Fig. 7b**). The T_h_ cells show a more active but less exhausted phenotype (**Fig. 7c**). Overall, these analyses confirmed that the Cd4^+^ T cells in the mouse heart VisiumHD datasets were indeed more functionally potent.

**Fig. 7.**
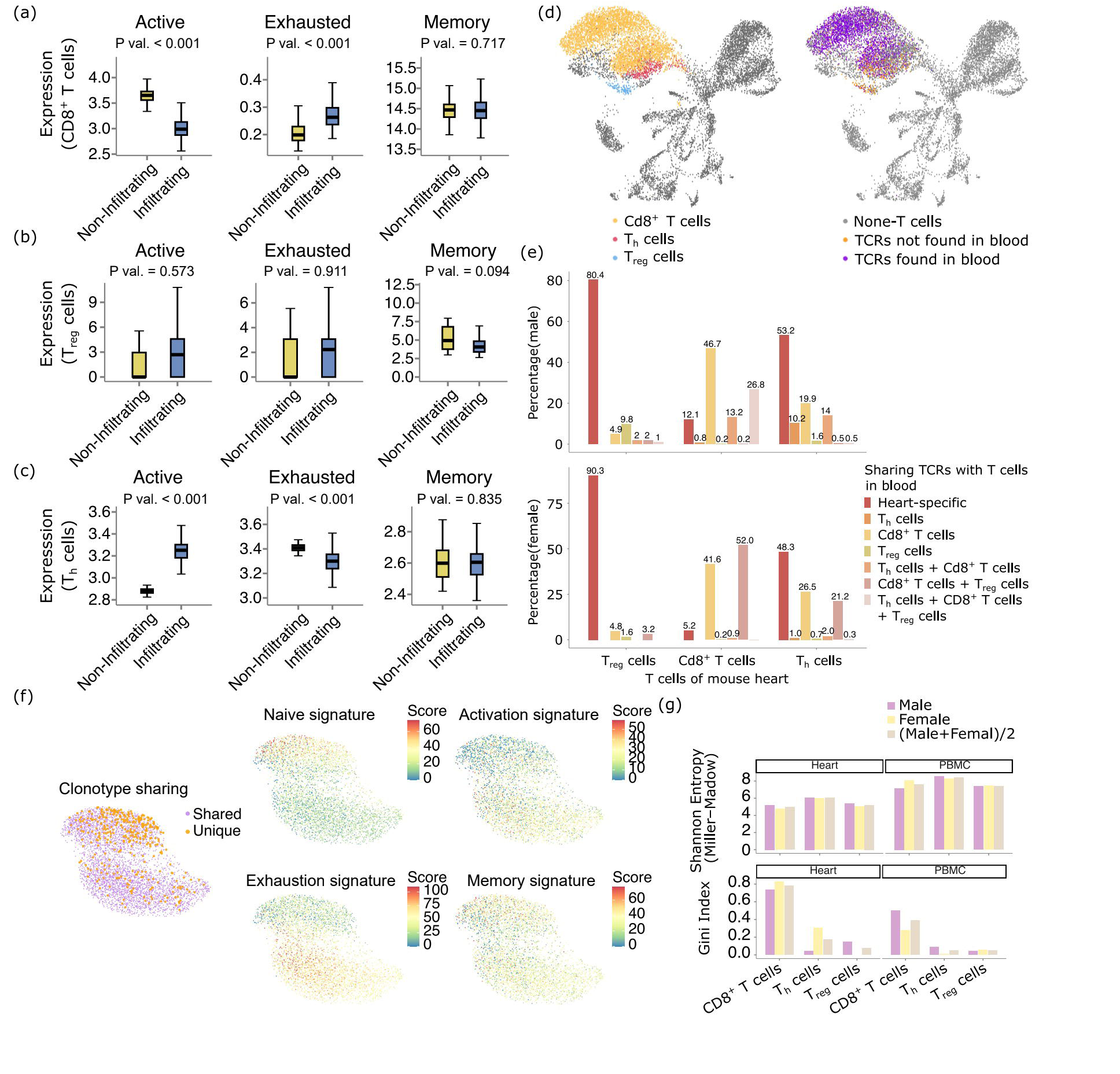
Functionally active CD4^+^ T cell responses in heart myocarditis. (a-c) The expression of activation, exhaustion and memory gene signatures in CD8^+^ T cells (a), T_reg_ cells (b), and T_h_ cells (c) in the mouse VisiumHD data. (d) UMAP of the scRNA-seq data in the heart of the male mouse. The cells were color-coded by the T cell subtypes and also by whether the T cells belong to TCR clonotypes that are unique to the heart or that are also found in the peripheral blood. (e) The percentages of the T cells in each T cell subtype that belong to TCR clonotypes that are unique to the heart or that are found in the peripheral blood. For the latter category, the T cell clonotypes in the peripheral blood are also labeled by the functional subtype(s) of the peripheral blood T cells in the clonotypes. (f) UMAP and clustering analyses focusing on the Cd8^+^ T cells. (g) Shannon entropy and Gini index metrics for the TCR clonotypes found in the heart and the peripheral blood, for the different T cell subtypes and also for both mice.

To offer finer granularity into the functional status of the cardiomyocyte-engaging T cells, we generated scRNA/TCR-seq data from the peripheral blood and heart of the *Pdcd1*^−/−^*Ctla4*^+/–^mice. In **Fig. 7d**, We showed a UMAP of all cells from the heart of the male mouse. On the left figure, we highlighted the different subtypes of T cells, and on the right figure, we highlighted whether the T cells carried TCRs that are found in the T cells in the blood. Overlaying the two figures together, we found that very high percentages of the T_reg_ and T_h_ cells are heart-specific (the leftmost red bars in each group in **Fig. 7e**). On the other hand, the Cd8^+^ T cells mostly have shared clonotypes with Cd8^+^ T cells from the blood, suggesting that a higher proportion of these Cd8^+^ T cells are likely to be by-standers. Judging from **Fig. 7e**, it is also obvious that the Cd8^+^ T cells form two separate clusters, while the heart-specific T cells are mostly concentrated in only one of the clusters. We separately performed UMAP and clustering analyses of the Cd8^+^ T cells in **Fig. 7f**, which again revealed two separate clusters. The heart-specific Cd8^+^ T cells appear in the top cluster, which is smaller. We examined the expression of these T cells (**Fig. 7f**), and found that the top cluster consists of T cells that are more naive, less activated and less exhausted. Finally, we also examined the clonality of the TCRs from the single cell sequencing data in both the mouse heart or blood. As is shown in **Fig. 7g**, compared with the blood, the TCR repertoires of all of Cd8^+^, T_reg_ and T_h_ cells overall demonstrated higher bias in clonality and stronger immuno-dominance ^3,38^ of a minority of clones in the heart. This is attested to by their lowered Shannon entropies and higher Gini indices ^39^.

Overall, results from spacer and also evidence from the scRNA/TCR-seq data both point to functionally active CD4^+^ T cell responses in heart myocarditis, which should be carefully studied. Importantly, prior descriptive spatial analyses could not reveal this insight, as they would focus on the CD8^+^ T cells that are more abundant in the inflamed hearts.

## Discussion

A fundamental challenge in biology is understanding the “postal code” problem: how do cells know where to go in complex tissues? We developed a fast, effective, and fully interpretable multi-instance learning neural network to excise the cellular recruitment and engagement signals from the transcriptomic and spatial modalities of the SRT data *via* an attention mechanism. While the field has moved towards creating (near) single cell resolution and WTX SRT technologies, analytical methods that can provide true novel mechanistic insights have been lacking. Researchers are still mostly using SRT data to visually assess the spatial distribution of the various types of cells in tissues and manually compare gene expression across physical locations. This is helpful for answering many biological questions, but is far from harnessing the full potential of SRT, especially given the very expensive costs of generating such data. In this study, our key conceptual argument is that cellular recruitment and engagement rules can be directly learned and interpreted from spatial patterns in native tissues. This inverts the experimental paradigm: rather than perturbing pathways one at a time, we extract generalizable principles by observing thousands of naturally occurring recruitment and engagement events *in situ*. This framework transforms SRT data from a descriptive map into a predictive blueprint for functional discovery. We advance the field by formalizing and tackling a fundamental question to answer the why and the how of the formation of tissue structures consisting of an admixture of cell types, based on analyses of SRT data. Because of our innovation, spacer has revealed novel biological insights that existing SRT data analysis methods cannot.

A central technical advance of this study is the development of spacer as an interpretable framework for elucidating the patterns of cellular engagement in tissues. A broad implication of this work lies in the growing movement toward interpretable AI for biology ^40^. Deep learning has been extensively applied to genomics data, yet most existing models are black boxes that provide strong predictive performance without offering insightful mechanistic understanding. Such methods often fall short in advancing biological discovery. Spacer addresses this critical gap by embedding biological principles directly into its neural network design. The bottom-up topology formulation of cellular recruitment as a multiple-instance learning problem reflects the biology of tissue structural organization, where the collective properties of local neighborhoods of resident recruiting cells govern the likelihood of an engaging cell’s presence. By modeling local structures as bags, spacer captures these multiple-to-one relationships in a principled way, thereby aligning the model’s logic with biological intuition. This design strikes a balance between predictive power and interpretability, distinguishes spacer from many prior black-box methodologies, and makes it a powerful biological discovery engine.

We argue that spacer has the capability of revealing novel biological insights and enabling translational applications that are not affordable by prior spatial data analysis tools. One of the many examples that we have shown in our results is the finding that CD4^+^ T cells are more responsive compared to CD8^+^ T cells during myocarditis. As another example, one implication of the findings from the application of spacer to human tumor SRT data is that spacer can potentially be used as a diagnostic tool for discovery of antigens that are targetable by T cells or B cells in each patient. As we have shown, many of the T cell-recruiting genes with top spacer scores encode known tumor antigens, or encode more antigenic peptides in their protein products than would be expected by random chance. Following upon our discoveries, these genes could be further confirmed for their immunogenicity and binding immune cell receptors could be identified *via* tetramer-type assays. Furthermore, we expect the cost of SRT to dramatically drop in the future, following what has been witnessed for Next Generation Sequencing (NGS) over the past two decades. If this is achieved, spacer can even be used for antigen discovery in each individual patient as an SRT-based diagnostic tool, enabling personalized therapies. Indeed, while many NGS-based diagnostic, prognostic and predictive biomarkers have been moved into the clinic already, little has been done for SRT, mainly due to its formidable cost. The recent development of optics-free spatial genomics technologies offers the hope for this possibility ^41^, which claims to cost approximately $30 per tissue section.

One caveat of our work is rooted in the inherent technical difficulties still faced by the high definition SRT itself, which is the lack of accurate cellular segmentation from the pathological imaging data. Most of the high definition SRT data we used in this study are from VisiumHD. It is a known issue to the field that the cellular segmentation of VisiumHD data is challenging, and many researchers have just resorted to the 8μm binned counts, as was done in this work as well. The obvious issue is that the resulting data are only “near” single cell resolution, and sometimes two neighboring cells that are in contact will contribute counts to the same bin, thus muddying the data. However, it is important to note that this issue is independent of spacer itself, as it takes whatever the preprocessed data users choose to feed it. In this work, we carefully performed cell typing and have discarded bins of ambiguous cell types, which should have alleviated this concern to a large extent.

## Materials and methods

### High level overview of spacer

Spacer is a biologically constrained multiple instance learning (MIL) neural network designed to model how local gene expression and spatial proximity of neighboring cells influence the positioning of specific cell types within tissues. For each center cell, it collects surrounding “recruiting” cells within a defined radius and uses their gene expression profiles and distances as input. The model includes three main modules: (1) a distance attention module that prioritizes biological proximity patterns; (2) a gene feature module that identifies influential expression patterns; (3) a gene-weighting module that infers specific attractive or repulsive potential of each gene expressed in the recruiting cells. Spacer is to be deployed to one SRT dataset or multiple SRT datasets jointly, and the core deep learning model is trained with the bags collected from the SRT data, to estimate the effect of how gene expression impacts cellular recruitment and engagement. Trained with binary cross-entropy loss and early stopping to ensure robust generalization. Spacer outputs both a probabilistic map of cellular engagement and interpretable gene-level “spacer scores” that rank genes by their contribution to cell recruitment. To prioritize robust biological signals, we filter for highly expressed genes across recruiting cells. As the selected genes are only a small subset of all genes captured by the SRT data, having WTX SRT data to start with becomes especially important for ensuring sufficient initial feature space for the spacer analyses. Even after filtering of genes as mentioned above, the model retains a rich and unbiased pool of candidate genes for high-precision discovery. A detailed description of spacer is provided in **Sup. File 1**.

### General considerations for SRT data preprocessing and annotation

Raw SRT data were processed with the Python Scanpy (v 1.1) package. The expression matrices were filtered to exclude low-quality barcodes (<100 detected genes). Counts were library-size normalized and log1p-transformed. Variance stabilization was performed with the SCTransform workflow in Seurat, with the object split by sample, if needed, prior to transformation (*e.g.*, Seurat_obj[[“RNA”]] <-split(Seurat_obj[[“RNA”]], f = Seurat_obj$sample)). A set of 5,000 highly variable genes (HVGs) was selected for subsequent steps. Principal component analysis (PCA) was run on scaled HVGs (30 principal components), and sample-specific batch effects were mitigated in PCA space using IntegrateLayers with method = “HarmonyIntegration” to apply Harmony-based integration. The integrated embeddings were used to construct a shared nearest-neighbor graph (FindNeighbors), followed by unsupervised Louvain clustering with FindClusters and non-linear dimensionality reduction with UMAP (RunUMAP).

For high-definition tumor datasets, we constructed curated gene signature panels for the major cell types, including tumor cells, T cells, B cells, macrophages, fibroblasts, and endothelial cells. For each panel, we computed the summed normalized expression of the corresponding marker genes detected in each dataset. These signature scores were projected onto the UMAP embedding, and Leiden clusters were assigned biological identities by considering both the signature enrichment patterns and their spatial organization. For the Slide-tag dataset, we adopted the cell type labels provided by the original authors. For low-definition SRT data, we first calculated gene signature scores for each cell type. Because individual spots in these platforms capture transcripts from multiple cells due to their larger physical size, signature values represent mixed contributions from heterogeneous populations. To distinguish enriched *vs.* background signals, we binarized the spots based on the 75% percentile of the corresponding signature distribution in each dataset to assign cell types to the spots. As a single sequencing spot in Visium captures cells of a mixture of several cell types, one spot could possibly be assigned to be positive for >=2 cell types. However, we would like to note that this does not interfere with the input for our model, either for the recruiting cells or the engaging cells. For the myocarditis dataset, we employed a ChatGPT-based approach ^42^ to perform initial cell type annotation, which was subsequently refined using a gene signature–based method.

### Human subjects

This study was performed in accordance with Institutional Review Board protocols at Baylor College of Medicine (BCM) and UT MD Anderson Cancer Center (MDACC). For the immunopeptidomics study (BCM H-47389), informed consent was obtained for the collection of clinical data and biospecimens. A prospectively maintained, single-institution database was retrospectively queried. Eligible patients were patients with unresectable or recurrent histologically confirmed PM who received single or combination checkpoint immunotherapy from 2015 to 2023. For the Xenium-TCR-seq study (BCM H-40168), head and neck squamous cell carcinoma (HNSCC) tumor specimens were obtained from patients treated at two Baylor College of Medicine-affiliated hospitals, Harris Health Ben Taub Hospital and the Michael E. DeBakey Veterans Affairs Medical Center. For the CosMX study (MDACC 2014-0938), patients with treatment-naïve, surgically resectable gastric cancer were evaluated at MDACC and underwent standard-of-care gastrectomy. Immediately after resection, specimens were grossed and sampled by an experienced pathology assistant under supervision of a gastrointestinal pathologist. Tumors were staged according to the AJCC criteria; the CosMx cohort comprised two gastric cancers (one pT3 tumor and one pT4 tumor). Tissue used for spatial transcriptomic profiling was obtained from banked or residual material under written informed consent in accordance with an institutional review board (IRB)–approved protocol. All procedures complied with the Declaration of Helsinki and relevant institutional and federal regulations governing research on human subjects.

### Immunopeptidomics *via* mass spectrometry

We used purified anti-human HLA-ABC pan antibody (W6/32) purchased from BioXcell and purified InVivoMAb anti-mouse MHC Class I antibody (H2-Kd: clone SF1.1.10, Bio X Cell Cat# BE0077) purchased from BioXcell. The tumor sample with lysis buffer was centrifuged at 20,000 G for 30 min at 4°C to isolate pMHC. Subsequently, 50 μl of anti-MHC-I antibody conjugated magnetic beads were added to the supernatant and incubated for 2hr at 4°C with gentle agitation. After incubation and meticulous washing, MHC molecules and peptides were eluted, segregated, and eventually subjected to a Bruker timsTOF Ultra2™ mass spectrometer. We extracted ligand peptides bound to MHC-I through immunoprecipitation and analyzed them using high-resolution MS. The acquired spectra are searched against the target-decoy human and mouse RefSeq database in the FragPipe computational platform with the no-digest enzyme. Assigned peptides were filtered with a 5% false discovery rate.

### Classification of human protein localization

The human_compartment_integrated_full.tsv file was downloaded from: compartments.jensenlab.org/Downloads ^43^. A protein can be located at multiple locations (multiple annotation entries for one protein). We only kept localization annotations with a confidence score>4. Within the remaining annotations, if a protein is documented as an extracellular protein for any entry, this protein would be classified as extracellular. For the remaining proteins, if a protein’s entries are all intracellular, this protein would be classified as intracellular. For proteins that are not classified as either extracellular or intracellular in the above two steps, their localization is recorded as uncertain. **Sup. Table 3** shows the classifications that we assigned to all proteins.

### Gastric cancer CosMX WTX data generation

Representative formalin-fixed, paraffin-embedded (FFPE) tumor blocks were selected by an experienced pathologist to include viable tumor, adjacent stroma, and interface with non-neoplastic mucosa where available. From each selected block, serial 5-µm sections were cut; one section was stained with hematoxylin and eosin (H&E) for histologic review and pathology-guided region selection, and adjacent sections were reserved for CosMx SMI processing. FFPE sections designated for spatial transcriptomics were mounted onto the center area (20mm x 15 mm) of Superfrost Plus Micro Slide (VWR, Cat# 48311-703) following the guide of CosMx Spatial Molecular Imager (SMI) RNA assay (Bruker Spatial Biology). Sections were processed using the CosMx Human Whole Transcriptome (WTX) RNA assay according to the manufacturer’s instructions. Tissue sections were placed within the capture area, and tiled fields of view (FOVs) were acquired to cover the majority of the tumor-containing region. Multimodal cell segmentation combined nuclear staining, morphology markers, and transcript-based refinement using the CosMx SMI/AtoMx Spatial Informatics Platform software to derive single-cell boundaries and segmentations. CosMx WTX gene count matrices and associated metadata were imported into R and processed using Seurat (v5.x). Cells with fewer than 20 detected genes were removed, and low-quality cells lacking expression of canonical lineage markers were excluded. Genes detected in only a small fraction of cells were also filtered out prior to normalization. After quality control, a high-quality set of single cells was retained for downstream analysis.

### Classification of tumor regions for the Xenium-TCR-seq data

The methodology of Xenium-TCR-seq was described in our preprint ^24^. Tumor cells within the Xenium-TCR-seq data were identified through Leiden clustering and differential gene expression analysis. Using the centroid location of each cell, the proportion of tumor *vs.* total cells within a radius was used to classify whether a cell was located in a tumor or non-tumor region. For the human samples 231120, 2309132, and 231004B2, since tumor cells consisted of the predominant portions of the samples, non-tumor or stromal region cells were defined as those with less than 50% tumor cells within a 30 μm radius, and all other cells were classified as located within tumor regions.

### Mouse myocarditis model

All animal experiments were performed in accordance with the guidelines and regulations of the Institutional Animal Care and Use Committees at Baylor College of Medicine (BCM) (AN-6685). We utilized *Ctla4*^+/-^*Pdcd1*^+/-^ mice on a C57BL/6J background, which were generously provided by Dr. James Allison at MD Anderson Cancer Center (Houston, TX) and his established breeding strategies were followed as previously described ^30^. For genotyping, genomic DNAs was extracted using DNeasy Blood & Tissue Kit (Qiagen, Cat# 69504), and PCR was performed with primers specific for *Ctla4* and *Pdcd1* according to previously published protocol. Mice exhibiting signs of distress, including lethargy, moribund appearance, respiratory difficulty, or failure to thrive, or as recommended by veterinary staff, were humanely euthanized using CO_2_ or inhalant anesthetic overdose, followed by bilateral thoracotomy.

### Mouse spatial transcriptomics workflow

Spatial transcriptomic profiling was performed using the Visium HD Spatial Gene Expression platform (10x Genomics), following the manufacturer’s protocol. Formalin-fixed paraffin-embedded (FFPE) mouse heart tissues were sectioned at 5 µm thickness and mounted on Visium HD slides. Standard hematoxylin and eosin (H&E) staining was performed, and slides were microscopically scanned. Then, spatial transcript capture was enabled by hybridizing tissues with the Visium Gene Expression Probe V2 using the CytAssist (10X Genomics). Probes targeting polyadenylated RNA were ligated and spatially transferred to the barcoded slide, followed by cDNA synthesis and library construction. Sequencing libraries were sequenced on the Illumina NovaSeq 6000 system using paired-end reads with dual indexing (43 cycles for Read 1T, 10 cycles for i7 Index, 10 cycles for i5 Index, and 50 cycles for Read 2S). Sequencing depth was determined by multiplying the percentage of the capture area covered by tissue (over 50%) by 275 million read pairs.

### Single-cell isolation from mouse heart tissues and blood

Briefly, mice were euthanized, and whole blood was obtained *via* cardiac puncture. After perfusion with PBS *via* the right ventricle, hearts were immediately harvested, minced, and enzymatically digested using a mouse tissue dissociation kit (Miltenyi Biotec Inc., Auburn, CA, USA, Cat# 130–096–730) at 37°C for 10 minutes in a rotating incubator. The digested mixture were sequentially filtered through 100 μm cell strainers (Corning Life Sciences Plastics, Cat# 431752), and red blood cells were lysed using ACK lysing buffer (ThermoFisher Scientific, Cat# A1049201), Cells were then washed with unsupplemented RPMI 1640 media (Corning, Cat# 15-040-CV). The cell pellets were resuspended and filtered again through 70 μm cell strainers (Corning Life Sciences Plastics, Cat# 431751), centrifuged, and washed with RPMI 1640 with L-glutamine (Corning, Cat# 10-040-CV) supplemented with 10% fetal bovine serum (FBS, GenDEPOT, Cat# F0910-050). Finally, cells were cryopreserved in a solution of 10% DMSO and 90% FBS at -80°C. For long-term storage, cells were placed in the vapor phase of a liquid nitrogen storage system.

Whole blood was collected into EDTA-treated tubes and immediately diluted 1:1 with Dulbecco’s phosphate-buffered saline (DPBS) without calcium and magnesium (Corning, Cat# 21-031-CV), supplemented with 2mM EDTA (Millipore, Cat# 4055-100ML). The diluted blood was gently layered over 4.5 mL of Ficoll-Paque (Cytiva, Cat# 17-5442-02) in a 15mL tube and centrifuged at 800 g for 20 minutes at 20°C without brake. The plasma was collected separately, and the peripheral blood mononuclear cells (PBMCs) were carefully transferred to a new tube. PBMCs were washed with DPBS/EDTA solution, treated with ACK lysis buffer, and washed again with DPBS. The cell pellet was then rinsed with unsupplemented RPMI 1640, followed by RPMI 1640 supplemented with 10% FBS. Cells were resuspended in FBS containing 10% DMSO and cryopreserved at -80°C. For long-term storage, samples were transferred to the vapor phase of a liquid nitrogen storage system.

### Single-cell library preparation and sequencing

Single-cell suspensions from each heart tissue and matched PBMC sample were loaded into separate wells of a Chromium Next GEM Chip N (10X Genomics, Cat# 2000418) to generate gel beads-in-emulsion (GEMs), each containing uniquely barcoded beads for downstream single-cell capture and indexing. Cell concentration was adjusted to approximately 700-2,000 cells/µL to target recovery of up to 20,000 cells per sample. Sequencing libraries for 5’ gene expression and T cell receptor (TCR) V(D)J profiling were prepared using the Chromium Next GEM Single Cell 5’ HT Reagent Kits v2 (10X Genomics, Cat# 1000356), the Mouse Chromium Single Cell V(D)J Amplification Kits (10X Genomics, Cat#1000254) and the Library Construction Kit (10X Genomics, Cat# 1000190) according to the manufacturer’s instructions. All libraries were sequenced on the Illumina NovaSeq 6000 system using paired-end reads with dual indexing 26 cycles for Read 1, 10 cycles for i7 Index, 10 cycles for i5 Index, and 90 cycles for Read 2), targeting 400 million reads per gene expression library and 100 million reads per TCR amplification library.

### Single-cell RNA sequencing data processing and data integration

Sequencing reads were aligned to the mouse reference transcriptome (mm10) using the Cell Ranger multi pipeline (10x Genomics, v9.0.0)^25^ with default parameters. This pipeline enables the joint processing of gene expression and TCR V(D)J profiling data from the same cellular libraries, allowing for the simultaneous quantification of transcriptomic profiles and immune receptor sequences. Scrublet was integrated within the Scanpy framework to identify and remove potential doublets ^44^. Stringent quality control criteria were applied, and cells were excluded if a cell expressed fewer than 100 total unique molecular identifiers (UMI) counts, or fewer than three genes, or zero or greater than 20% of total UMI of mitochondrial genes. Normalization and batch correction were implemented using scVI *via* scvi-tools ^45^. Sample ID was used as the batch variable to correct for inter-sample variability, ensuring robust data integration.

### Statistical analyses

Pathway enrichment analyses were performed using GOrilla ^6^. Trend test was performed by the jonckheere.test function from the clinfun R package (1.1.5). For gene network analysis, we used hdWGCNA (v0.4.05) ^21^. Gene set enrichment analysis (GSEA) was conducted with clusterProfiler (4.14.6) ^46^. For the boxplot, box boundaries represent interquartile ranges, whiskers extend to the most extreme data point, which is no more than 1.5 times the interquartile range, and the line in the middle of the box represents the median. All statistical tests are two-way, except for **Fig. 3e-h**, where a clear directional alternative hypothesis is present.

## Acknowledgments

None

## Author contributions

J.Y., J.Z.,W.C. contributed to the bioinformatics analyses. J.S., S.K., J.C.,Cl.L.,H.J.,B.B., H.L. contributed to the generation of the human tumor immunopeptidomics data and the mouse VisiumHD and immunopeptidomics data. K.M., W.H. contributed to the generation of the Xenium-TCR-seq data. Ch.L., H.L., T.W. contributed to funding acquisition. W.H., T.Y., H.L. contributed critical insights to the study. Y.L., I.L., L.S., S.H., K.C., K.K., P.M., L. We., L.Wa. contributed to the gastric cancer CosMx data generation. K.W., H.Z. contributed to validation experiments for the human tumor spacer scores. T.W. supervised the overall study.

## Data and code availability

The public human Visium and VisiumHD data are available at 10X Genomics’s website (accession IDs shown in **Sup. Table 1**). The Slide-tag dataset is available from ^1^. ***<u>The gastric</u> <u>cancer CosMx data that we generated will be made available upon manuscript</u> <u>acceptance.</u>*** Our Xenium-TCR-seq data are available from GSE300147. The mouse heart VisiumHD data that we generated are available at: GSE298932. The mouse scRNA-seq/TCR-seq data that we generated are available at GSE299032. ***<u>The human and mouse immuno-peptidomics data will be made available upon manuscript acceptance.</u>*** The bulk RNA-seq data across developmental stages (fetal, child and adult) were obtained from the Temporal Expression during Development Database (TEDD) 2.0 ^18^. Data for the ICI-treated cohorts are available through their original publications ^13–16^. Spacer is available at: spacer-readme.readthedocs.io.

**Sup. Fig. 1** Tumor cell-expressed genes that repel T cell recruitment are related to the tumor extracellular matrices. Showing GO analysis of the ranked genes from **Fig. 3a**.

**Sup. Fig. 2** Shared and unique recruitment mechanisms across tissue types and engaging cell types. (a) Overlaying the H&E staining image with T cells defined from the VisiumHD gene expression data for the non-malignant kidney sample. (b) Showing only the T cells defined from the VisiumHD gene expression data for the non-malignant kidney sample. (c,d) GO analyses of the top tumor cell-expressed genes predicted by spacer to promote stromal/immune cell infiltrations. Showing results for fibroblasts (c) and endothelial cells (d) in this figure.

**Sup. Fig. 3** Advantages of SRT in yielding insights on cellular infiltration into tissues. Showing Spearman correlations between the bulk RNA-seq expression of the genes that are ranked by spacer as the top genes with the strongest T cell recruitment potentials and the predicted infiltration levels of CD8^+^ and CD4^+^ T cells calculated based on a signature approach applied to the bulk RNA-seq data. (a,b) CD8^+^ T cells, (c,d) CD4^+^ T cells. (a,c) show the histogram aggregating all individual correlation coefficients in (b,d).

**Sup. Fig. 4** Xenium-TCR-sequencing data. (a) UMAP based on the expression of the cells from the Xenium-TCR-seq data. (b) The different types of cells shown in their spatial contexts in the Xenium-TCR-seq data. (c) The most expanded TCR clonotypes in the Xenium-TCR-seq data.

**Sup. Fig. 5** GSEA analyses for testing the enrichment of T cell, B cell and macrophage genes that are more highly expressed in the “predicted not to infiltrate” regions than the “predicted to infiltrate” in the mitochondrial components that are not related to the TCA cycle.

**Sup. Table 1** The SRT datasets involved in this study

**Sup. Table 2** The spacer scores for all engaging cell types involved in the human tumor and mouse heart datasets

**Sup. Table 3** The cellular localization classifications assigned to all human proteins

**Sup. Table 4** Detailed results for the differential gene expression analyses of T cell migration-and stemness-related genes between the “predicted to infiltrate” regions and the “predicted not to infiltrate” regions. Negative values indicated higher expression in the “predicted not to infiltrate” regions.

**Sup. File 1** Detailed model description

**Sup. File 2** Additional results and discussion

